# Mapping the impact of age and APOE risk factors for late onset Alzheimer’s disease on long range brain connections through multiscale bundle analysis

**DOI:** 10.1101/2024.06.24.599407

**Authors:** Jacques Stout, Robert J Anderson, Ali Mahzarnia, Zay Han, Kate Beck, Jeffrey Browndyke, Kim Johnson, Richard J O’Brien, Alexandra Badea

## Abstract

Alzheimer’s disease currently has no cure and is usually detected too late for interventions to be effective. In this study we have focused on cognitively normal subjects to study the impact of risk factors on their long-range brain connections. To detect vulnerable connections, we devised a multiscale, hierarchical method for spatial clustering of the whole brain tractogram and examined the impact of age and APOE allelic variation on cognitive abilities and bundle properties including texture e.g., mean fractional anisotropy, variability, and geometric properties including streamline length, volume, and shape, as well as asymmetry. We found that the third level subdivision in the bundle hierarchy provided the most sensitive ability to detect age and genotype differences associated with risk factors. Our results indicate that frontal bundles were a major age predictor, while the occipital cortex and cerebellar connections were important risk predictors that were heavily genotype dependent, and showed accelerated decline in fractional anisotropy, shape similarity, and increased asymmetry. Cognitive metrics related to olfactory memory were mapped to bundles, providing possible early markers of neurodegeneration. In addition, physiological metrics such as diastolic blood pressure were associated with changes in white matter tracts. Our novel method for a data driven analysis of sensitive changes in tractography may differentiate populations at risk for AD and isolate specific vulnerable networks.

## Introduction

Alzheimer’s disease (AD) causes progressive neurodegeneration and brain pathology associated with amyloid and tau proteinopathies, yet the etiology of the disease is not fully clear. Early onset AD patients, patients diagnosed under the age of 65, account for 5% of the patients, while late onset AD (LOAD) accounts for 95% of cases. Overall, age is the most prominent risk factor for LOAD; however, the APOE4 allele is a known major genetic risk factor. Thus it is imperative to understand how age and the APOE4 allele contribute to the neural correlates of vulnerability to AD.

Brain morphometry has established that vulnerable brain regions (e.g. entorhinal cortex, hippocampus), decline early on, but less is known about potential early changes in the structural connection of these regions and their relative timelines. Diffusion weighted imaging and associated tractography models can provide us with a better understanding of the dynamics of brain connectivity, specifically how structural connectivity is impacted by the integrity of micro-structural fiber bundles. It is unknown how the dynamics of these bundle properties at an individual’s level contribute to their disease risk profiles. Tractography studies have uncovered micro-structural changes in fiber bundle characteristics in Parkinson’s disease [1, 2], epilepsy [3, 4], major depression [5, 6] and AD itself [7–10]. Using whole brain diffusion weighted tractography analyses on individuals with the apolipoprotein (APOE) genetic risk at different points in their lives, one can determine how long white matter fiber bundle connections change in individuals at greater genetic risk (i.e., APOE4 allele carriers) and provide insight into disease propagation as well as help monitor subtle changes in response to disease or interventions.

It is known that structural brain changes may be detected decades before AD symptoms appear [11–14]. Additionally, several studies have demonstrated that cognitive measures from the National Alzheimer’s Disease Coordinating Center (NACC) Uniform Dataset (USD3) can be used to estimate the risk of developing LOAD before the condition is fully manifest [15, 16]. These tests include assessments for various cognitive, behavioral, and functional changes. In this study of cognitively normal subjects, we employed a subset of the standardized UDS3 cognitive measures, as well as sensory markers for visual attention, markers for olfactory memory, and additionnal physiological metrics, all to better put neurodegeneration in the context of brain-body relationships and cognitive decline.

Combining diffusion-weighted imaging tractography and UDS cognitive assessment data, we have conducted brain-wide white matter fiber bundle analyses with the aim of determining which specific connections differentiate cognitively normal APOE4 carriers from non-APOE4 individuals. Further, we sought to determine if any fiber bundle connections can be linked to either age, genotype, or cognitive abilities in cognitively unimpaired persons.

While connectome analysis can elucidate correlations between genotypes and specific white matter fiber connections, the magnitude of multiple comparison statistical correction is so great that few findings survive. To address this issue, we used a multi-level, data-based tractography clustering approach to identify bundles of interest, rather than a label-based approach, iterating on previous tools for fiber bundle splitting and analyses [17–19]. Our agnostic data driven, whole brain analytical approach allows for the identification of major coherent fiber tract paths and relate those fiber bundle characteristics to subject meta data.

We examined diffusion-weighted tractography characteristics from ages 20 to 80 to observe age and genotype specific changes in individuals at heightened genetic risk for LOAD (i.e. enriched for APOE4), relative to non-APOE4 carrier controls. Our aims were: 1) to determine which white matter fiber bundle metrics may be sensitive to age-related decline and genotypic risk for LOAD; 2) to assess for possible relationships between fiber bundle properties and cognitive performance, and the interaction of age by APOE4 genotype carrier status; and 3) to improve the specificity of fiber mapping to determine vulnerable white matter bundles in a data driven, multi-level spatial clustering approach.

## Methods

### Subjects

All experimental procedures have been previously reported in (Mahzarnia et al, 2023), with this paragraph serving as an updated brief overview. The procedures were approved by the Duke University Medical Center Institutional Review Board. All participants, and where applicable study partners or caregivers, provided informed consent, and were compensated for their time at the end of the study. A total of 78 subjects were recruited, 33 males, and 45 females with a combined age range from 20 to 80 years (yrs) (median age 50.44 yrs, mean ± std: 50.4 ± 15.15 yrs). APOE genotype allelic distribution was 7 APOE2/APOE3 carriers (9% of study cohort); 35 APOE3 homozygous carriers (45%); 29 APOE3/4 carriers (37%), 7 APOE4/4 homozygous carriers (9%). Twenty-one participants (27%) had no known family history of LOAD and 57 (73%) had family relatives with LOAD.

### Physiological, Behavioral & Cognitive Evaluation

We measured body, mass, height, body mass index (BMI) and blood pressure as physiological parameters for subsequent analysis. Cognitive evaluation was based upon performances with the NACC UDS3 neuropsychological assessment battery [20]. The UDS3 tests used in this study included the Montreal Cognitive Assessment (MOCA) screen; Craft Story to assess unstructured, auditory-verbal narrative learning and memory [21]; the Trail Making Test, with parts A and B used to assess graphomotor and visual scanning speed, and part B assessing the complex task rule maintenance and switching under timed conditions [22]; Wechsler Adult Intelligence Scale – 3^rd^ revision (WAIS-3) Digit Symbol subtest to assess information processing speed [23]; Categorical Verbal Fluency (animals, vegetables) and Lexical Verbal Fluency (F-words, L-words), assessing speed of distributed lexical search and semantic memory [24] and the Digit Span subtest from the WAIS-3, assessing auditory verbal attention, as well as working memory when using the backwards test [25]. On top of the UDS3 battery of cognitive tests, we administered the Rey Auditory Verbal Learning Test (RAVLT) for assessing structured, auditory verbal word list learning and memory [26]

Additionally, we assessed olfactory sensory function, as well as short- and long-term memory for olfactory stimuli, measuring valence on aspects of odor pleasantness, intensity and familiarity [8]. Dynamic visuo-spatial attention abilities were assessed using the computerized Useful Field of View (UFoV) Test from BrainHQ [27] with both the presence of passive and active distractors [8]. For statistical analysis, all subject traits, and cognitive tests metrics were converted to a z-score and combined across cognitive domains as seen in Table 1.

**Table 1:** Summary of combination of different z-scores of cognitive tests into cognitive assessment levels.

| **Average measurement** | **Zscore of cognition test** |
| --- | --- |
| Olfactory Memory | Composite Familiarity, Composite Nameability, Number of odors recognized out of 6 |
| Verbal Memory | RAVLT Trial6, RAVLT Trial7, RAVLT LEARNING, RAVLT FORGETTING, RAVLT IMMEDIATE |
| Logical Memory | Story Immediate verbatim, Story Immediate paraphrase, Story Delayed verbatim, Story Delayed paraphrase |
| Verbal short-term Memory | Digit Span: forward total correct, Digit Span: forward max length |
| Working Memory | Digit Span: backwards total correct, Digit Span backwards max length, Trail Test: TrailB |
| Verbal Fluency | Fluency categorical, Fluency letters |
| Visual Attention | Trail Test: Trail A |
| Visuo Spatial | Test 2 of field of view, passive distractors, Test 3 of field of view, active distractors |

### Imaging

Magnetic resonance imaging (MRI) was performed using a 3T GE Premier Performance scanner with Signa Premier version (ver. 28) with a 60 cm gradient coil with peak strength of 115 mT/m, and a 48-channel head coil. Diffusion weighted images were acquired using a robust motion-resistant, multi-shot echo planar imaging (EPI) sequence with 4 interleaved shots and reconstructed using MUSE [28]. Parameters were set at TE 60.6 ms/TR = 14,725 ms, resolution of 1×1×1 mm isotropic, FOV at 25.6×25.6cm (256×256 matrix) and 136 slices. Overall, 21 diffusion directions (*b* = 1000 s/mm^2^) and 3 non-diffusion weighted images were acquired in 24 minutes. In addition, anatomical data were acquired using a 3D T1-weighted IR-FSPGR-BRAVO sequence with TE = 3.2 ms, TR = 2263.7 ms, flip angle 8°, prep time 900 ms, recovery time 700 ms, 256 x 256 x 160 slices, 25.6 x 25 x 6 x 16 mm FOV, with image acquisition time of 5 min, 17 s.

### Image and Bundle Analysis

All diffusion images were preprocessed using a pipeline [29, 30] that included FSL-BET[31] for brain masking, ANTs for co-registration and eddy current correction[32], and a Principal Component Analysis denoising process[33, 34]. The MRtrix toolbox [35] was used for creating diffusion parametric maps, (e.g., fractional anisotropy, FA) which provide important estimates of microstructural tissue integrity and additionnal voxel level diffusion characteristics [36].

The MRtrix toolbox [35] generated the streamlines of individuals using Anatomically-Constrained Tractography (ACT), in which diffusion maps were co-registered with subject-specific anatomical T1-weighted imaging data segmented into five tissue type maps (e.g., grey matter, sub-cortical grey matter, white matter, CSF and pathological tissue maps). This ACT correction method helps guarantee that generated streamline models are properly aligned with the individual brain anatomy of the subject [37], while also allowing anatomically truncated tracks to be re-tracked. White matter tract fiber streamline generation was carried out via a probabilistic method second-order integration of Fiber Orientation Distribution (FOD) [38] with grey matter seeding. A FOD cutoff of 0.06 was used with a step size of 0.1 mm, a minimum length of 2 mm, a maximum length of 250 mm, and a 45° limitation angle. Two million streamlines were generated per participant. The resulting streamlines were then registered into study-specific cohort average brain space using the same rigid and affine transformations applied via the ‘dipy’ package [34] and the resulting warp transforms were applied via mrtrix3 and the tcktransform function [39].

After ACT streamline quality control procedures, streamlines were subdivided across all participants into anatomically consistent and equivalent groupings based on pathing results. The multi-level spatial clustering approach was conducted by first downsampling streamlines downsampled into a set of 20,000 streamlines. Then, twenty individuals were randomly selected to be used as a standard template. The downsampled streamlines from all these individuals were combined into a single set, for which the right-side streamlines were flipped onto the left side of the brain to include input from both sides of the brain. From there, we formed our basic bundle centroids using dipy to form the basis of our future sets of bundles, that were made to be symmetrically equivalent for the left and right side of the brain, at first six pairs of centroids overall. The centroids were then compared to the streamlines of all subjects and used to extract the streamlines closet to the established centroids. Once this was done once initially, the process was repeated onto the already separated bundles, now establishing on every iteration three bundle descendants based on the previous set. This iterative process separates all streamlines of the brain into a set of 6, 18, 54, 162 pairs of ever smaller bundles that are equivalent across all subjects. This method allows us to divide our brain for all subjects into small sets of bundles that are 100% streamline based rather than based on previously established structures [17, 40].

From each bundle, we extracted multiple metrics, specifically the mean of the FA among streamlines, the standard deviation of the FA, the number of streamlines [41], the length of streamlines, the volume of the bundle (determined by the number of voxels traversed by the streamlines) [42–45], the asymmetry, L2 norm of FA measures, as well as the BUndle ANalytics for comparing left and right side bundles in order to determine their similarity in shape in each subject (after being superposed to each other) [46].

Once these statistics were extracted and summarized, the statistics for each bundle were compared to the age and genotype of subjects, using a linear model for most metrics, but using a quadratic model for average FA vs age [47, 48]. The bundle statistics were also compared to average cognitive measurements (see Table 1) and physiological data (Height, Weight, Systolic pressure, Diastolic pressure). The results were corrected with False Discovery rate (FDR) based on the number of bundles and the number of metrics each statistic was compared to. Our analysis process is illustrated in Figure 1.

**Figure 1.**
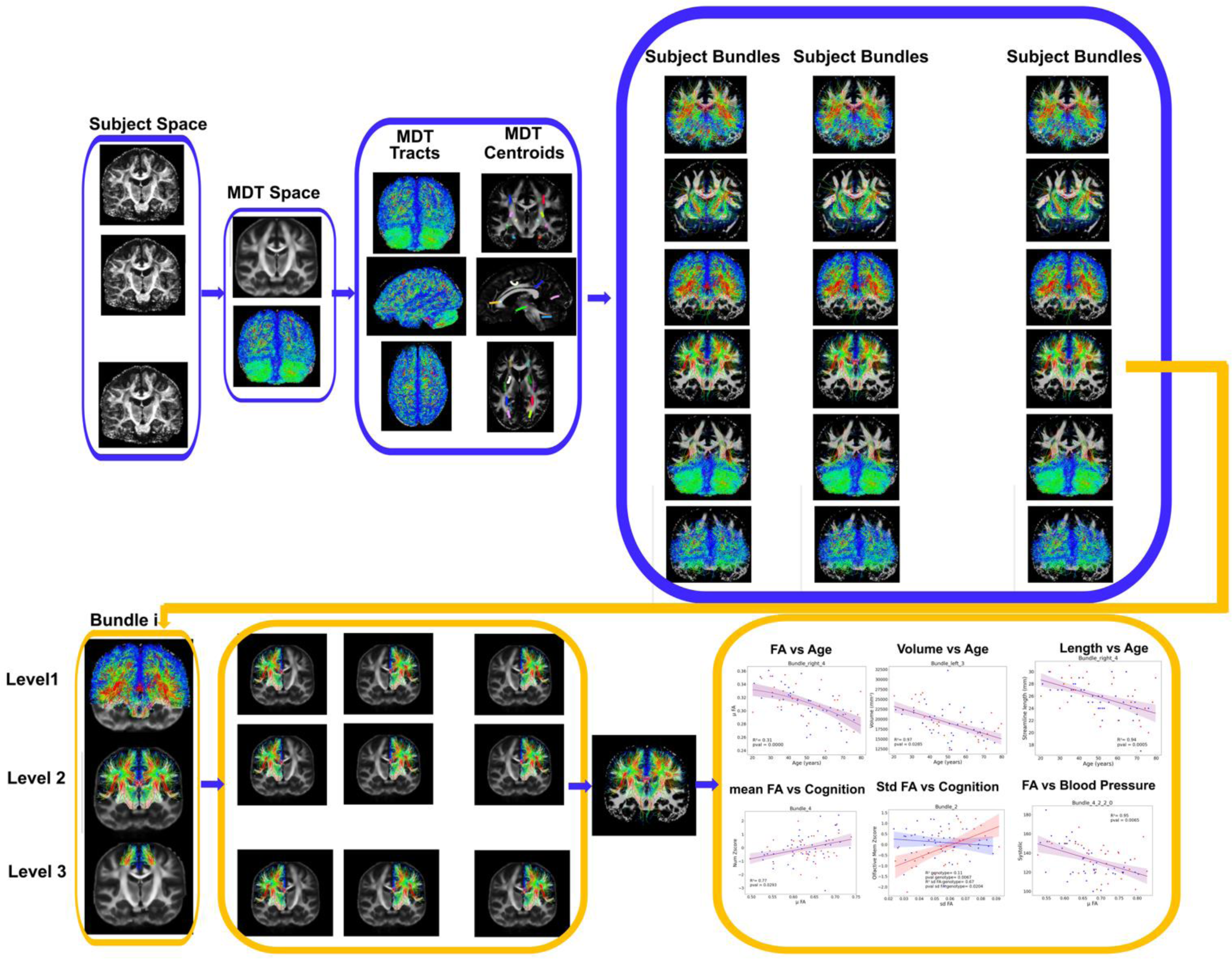
Flowchart of the approach. Diffusion weighted images are used to produce fractional anisotropy (FA) maps, registered into a mean deformation template (MDT) space. The calculated transforms are applied to subject tractograms to map them into the same space, before being clustered into four levels comprising 12, 36, 72, 144 bundles. The bundles are then characterized by texture properties reflecting diffusion attributes (e.g. mean FA and variance), and for volume, length, shape related metrics that are then used to test the impact of risk factors (age, APOE genotype, and their interactions), as well as to map how physiological and cognitive parameters impacted bundles.

## Results

### Multiscale brain tractography bundling

To tune our analyses to the scale of the effects impacting tracts along the aging trajectory we created a hierarchical representation of the brain tractography, spanning 4 levels. We parcellated the brain tractography in an iterative multiscale fashion based on spatial clustering (Figure 2), generating 6 sub-bundles per hemisphere at the first level, splitting each bundle in 3 sub-bundles at the next iteration, and so on. This allowed us to examine genotype, age, and genotype by age interactions in subjects at risk for late onset Alzheimer Disease (LOAD). We next examined the age, genotype, and age x genotype effects on bundle properties and the associations with cognitive traits at each level.

**Figure 2.**
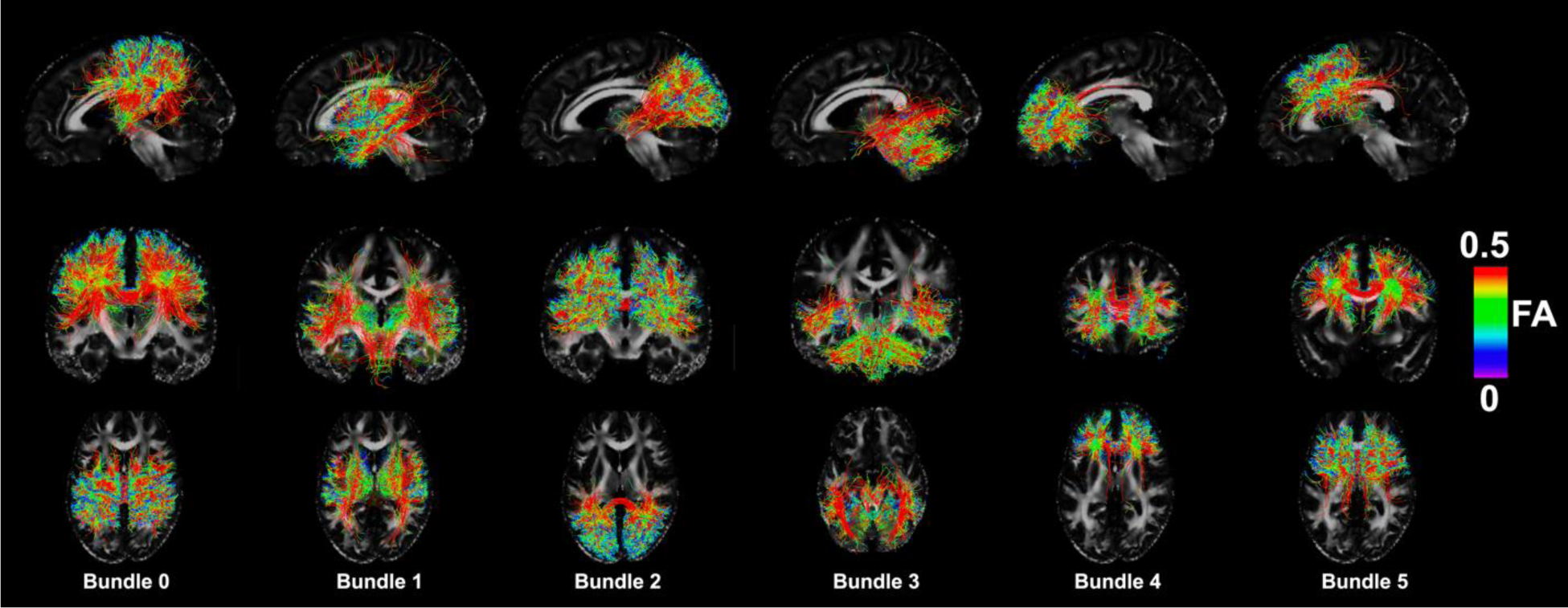
The first level of the multilevel clustering scheme shows the spatial distribution of the 12 bundles (6 on left side and 6 on right side).

**Table 2:** The anatomical regions corresponding to the top 6 bundles in one hemisphere at level 1 clustering.

| Bundle Id | Top Structures |
| --- | --- |
| Bundle 0 | Supramarginal - 8.14 % superior parietal - 5.65 % postcentral - 5.18 % |
| Bundle 1 | Superior temporal - 12.85 % thalamus proper - 12.42 % putamen - 6.97 % |
| Bundle 2 | Inferior parietal - 17.55 % lateral occipital - 10.74 % superior parietal - 9.56 % |
| Bundle 3 | Cerebellum cortex - 28.22 % fusiform - 3.88 % hippocampus - 3.85 % |
| Bundle 4 | Rostral middle frontal - 19.65 % Superior frontal - 10.87 % caudate - 4.21 % |
| Bundle 5 | Superior frontal - 15.57 % Caudal middle frontal - 8.21 % precentral - 7.48 % |

### Bundles with significant age effects

#### Significant FA effects

Looking at the effect on age on bundles, we identified a robust effect across the brain where the mean FA (Figure 3A) and variance of FA (Figure 3B) across the frontal lobe decreases with age, using a quadratic model for the relationship between mean FA and age [48–50]. While this effect could be seen across the brain, it was particularly significant in the bundles associated with the frontal lobe, and specifically such anatomical regions as the rostral middle frontal, superior frontal region, caudate nucleus and corticostriatal tract (See Supplementary Figure 1) as well as other frontal regions and connections. (See Table 3). The correlation of the FA variance with age tended to be higher than that of the mean FA even as it affected similar regions (Additional results can be found in Supplementary Figure 2 & 3).

**Figure 3.**
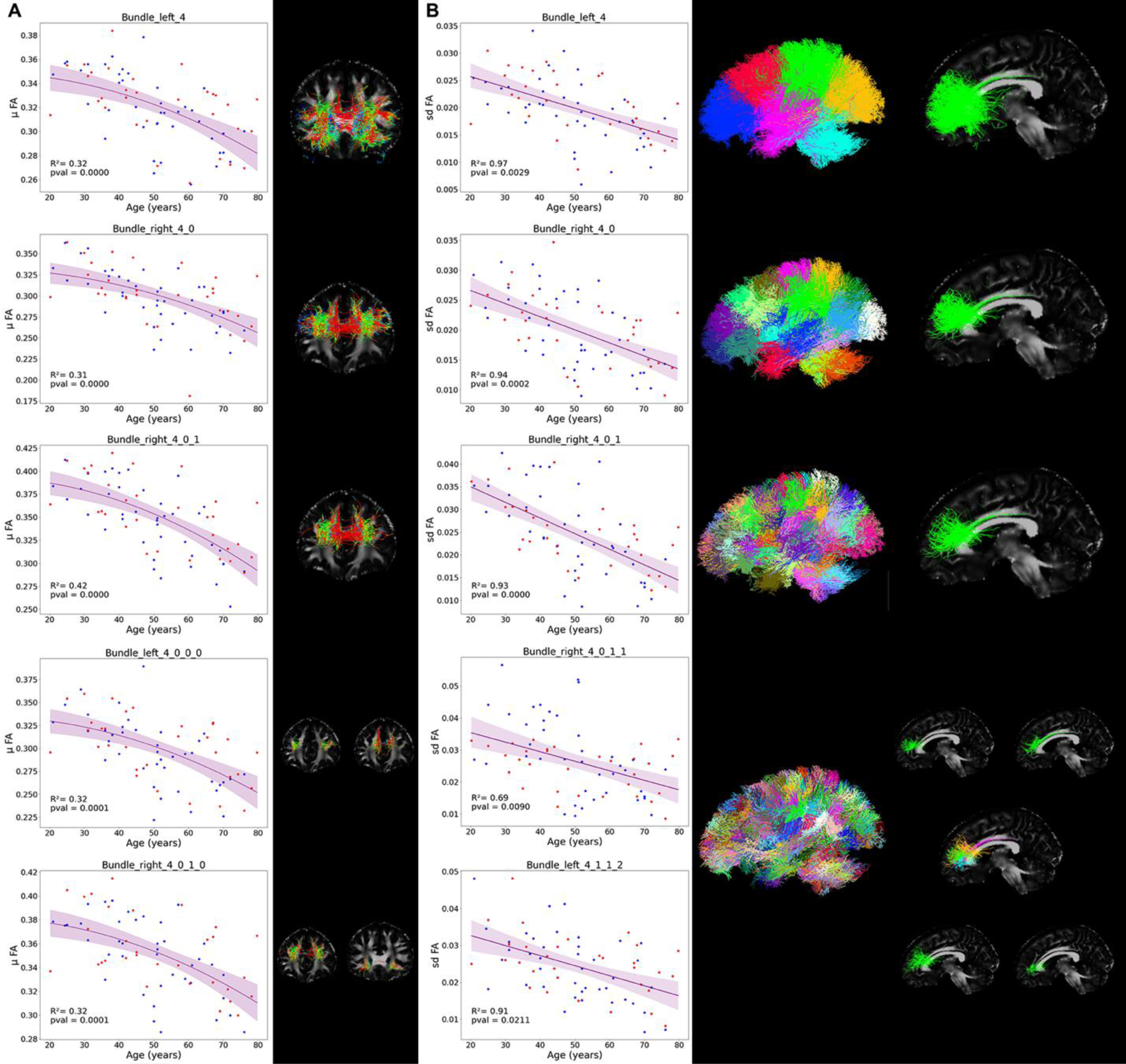
Significant bundle FA changes with age were focused on the frontal lobe for both A) average FA and B) variance of FA

**Table 3:**
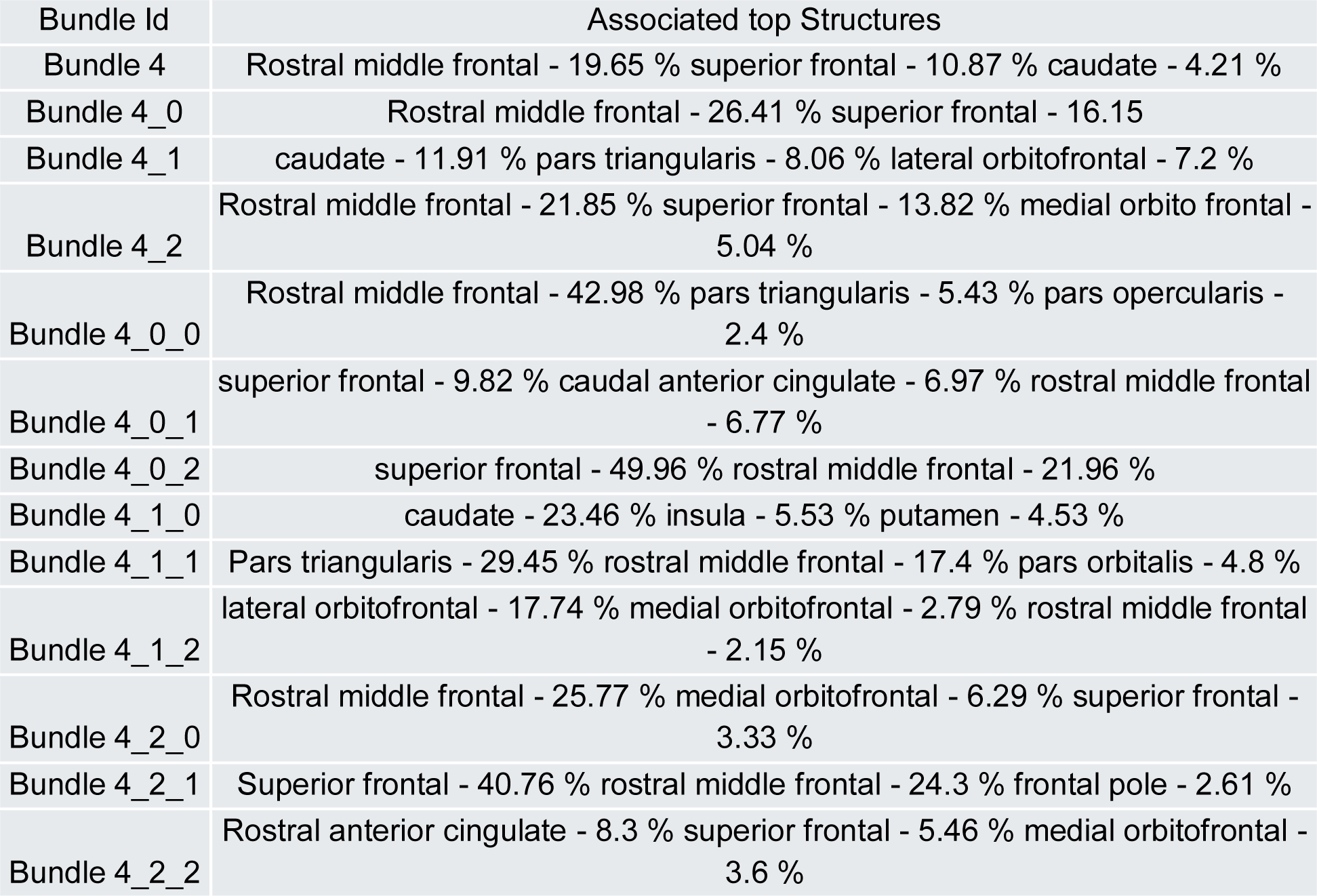
Correspondence between Bundle ID and top associated structures for frontal lobe bundles showing high correlation between average and variance of FA with age.

#### Significant age effects on bundle shape

While other metrics related to bundle shape and density were less affected by age overall, there was a significant effect in specific regions remaining after FDR correction. The volume of bundle streamlines was partially correlated with the number and length of streamlines, while also representing the overall spread of streamlines in a specific region. The volume of bundle streamlines decreased with age, with the effect being particularly visible in the cerebellum bundles, as well as one particularly associated with the supramarginal gyrus, another associated with the temporal to insula paths, and finally one associated with the superior parietal region and precuneus (Figure 4A).

**Figure 4.**
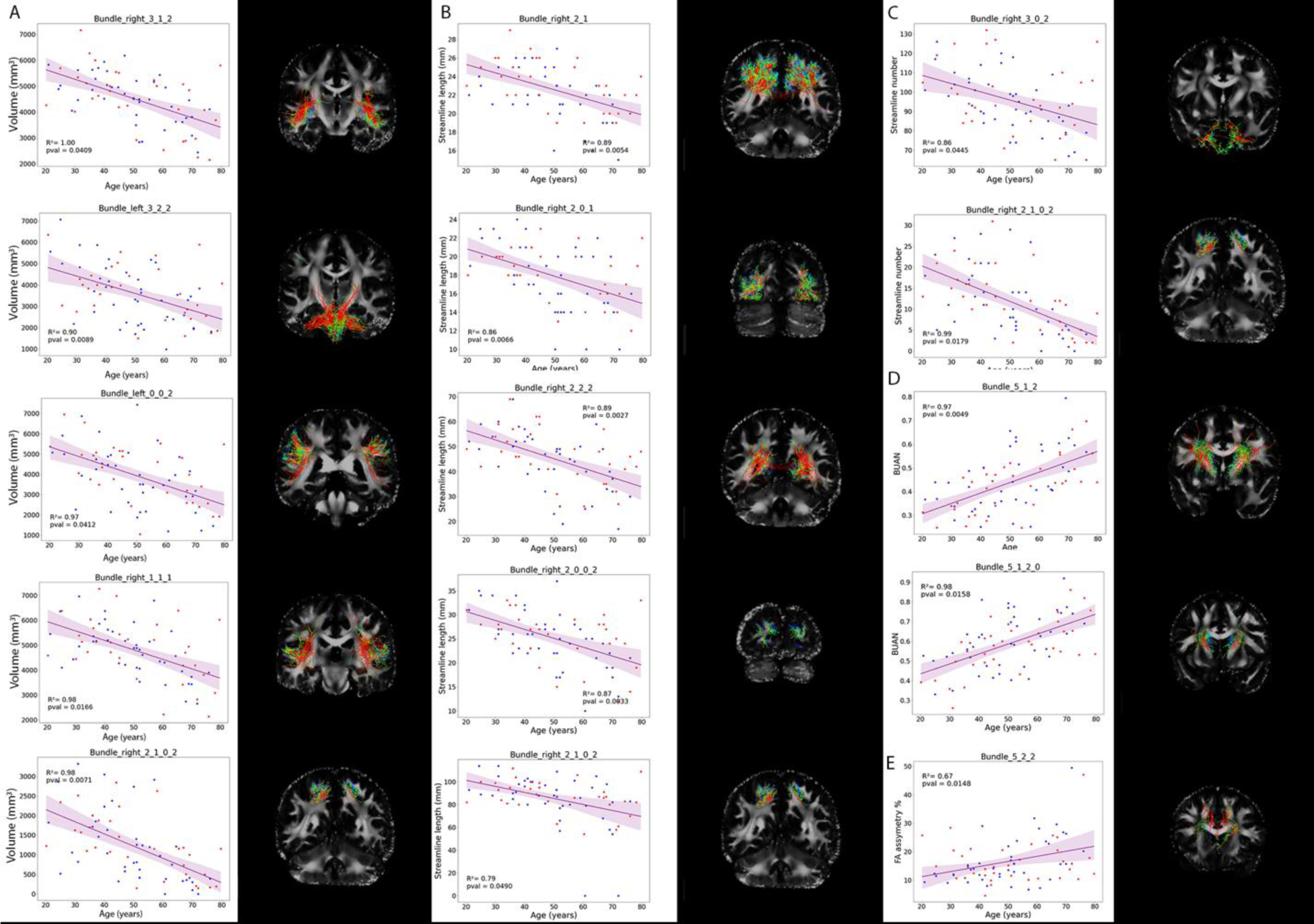
Significant changes with age in bundle properties for A) volume of bundle; B) length of streamlines; C) number of streamlines; D) symmetry of bundle shape; E) FA asymmetry of streamlines.

**Table 4:** Correspondence between Bundle ID and top associated structures for bundles showing high correlation between volume of bundle, number of streamlines or average length of streamlines, and Age.

|  |  |
| --- | --- |
| Bundle 3 | Cerebellum cortex - 28.22 % fusiform - 3.88 % - hippocampus - 3.85 % |
| Bundle 3_1_2 | fusiform - 5.65 % hippocampus - 4.02 % superior temporal - 1.25 %<br>*inferior longitudinal fasciculus* |
| Bundle 3_2_2 | Cerebellum cortex - 2.65 %<br>*cerebellar peduncles* |
| Bundle 0_0_2 | supramarginal - 45.69 % superior temporal - 15.68 %<br>*Middle longitudinal fasciculus* |
| Bundle 1_1_1 | Superior temporal - 19.93 % transverse temporal - 9.91 % insula - 5.26 %<br>*superior temporal sulcus, inferior & middle longitudinal fasciculus* |
| Bundle 2_1 | Superior parietal - 25.81 % inferior parietal - 20.72 % precuneus - 12.64 %<br>*superior parietal lobe, posterior region* |
| Bundle 2_1_0_2 | Superior parietal - 72.04 % precuneus - 14.31 % inferior parietal - 2.35 %<br>*superior parietal cortex, lateral posterior* |
| Bundle 2_0_1 | Lateral occipital - 46.56 % - inferior parietal - 10.77 % - fusiform - 10.48 %<br>*Occipital lobe lateral cortex* |
| Bundle 2_2_2 | Lingual - 7.94 % - precuneus - 2.26 % - pericalcarine - 1.16 %<br>*Inferior longitudinal fasciculus* |
| Bundle 2_0_0_2 | Lateral occipital - 49.04 % pericalcarine - 5.5 % - cuneus - 1.29 %<br>*Occipital lobe posterior cortex* |
| Bundle 2_1_0_2 | Superior parietal - 72.04 % precuneus - 14.31 % - inferior parietal - 2.35 %<br>*Superior parietal posterior cortex* |
| Bundle 3_0_2 | Cerebellum cortex - 45.51 %<br>*Inferior mid-lateral cerebellar cortex* |
| Bundle 2_1_0_2 | Superior parietal - 72.04 % precuneus - 14.31 %<br>Superior parietal cortex, posterior |
| Bundle 5_1_2 | caudate - 12.2 % thalamus proper - 3.59 % putamen - 2.06 %<br>*Insular region streamlines* |
| Bundle 5_1_2_0 | caudate - 21.55 % thalamus proper - 3.19 % putamen - 1.78 %<br>*External capsule* |
| Bundle 5_2_2 | Caudal anterior cingulate - 1.37 % thalamus proper - 1.36 %<br>*Superior longitudinal fasciculus* |

When comparing the length of streamlines in different bundles to age, we found a significant decrease in the length of streamlines with age for bundle streamlines that belonged to the superior parietal region, the lateral occipital cortex, and the lingual region (Figure 4B).

Very few bundles had their number of streamlines significantly impacted by age, however those that did had a negative correlation. These bundles corresponded to the frontal lateral cerebellum bundle and the superior parietal cortex (Figure 4C). We noted that the superior parietal cortex bundle had significant reductions in number of streamlines, length of streamlines, and overall volume reduction with age (with the latter having the highest correlation). The BUAN metric identified the level of shape similarity between the left and right symmetrical bundles in a specific subject. Two bundles showed an increase in BUAN with age, with one being the descendent of the other identifying that the caudate associated streamlines (and to a lesser degree those associated with the thalamus and putamen) had a significant increase in symmetry with age (Figure 4D), denoting possible loss of lateralization. Additionally, we observed that the bundle of streamlines situated above the corpus callosum bordering the caudate and cingulate have an FA asymmetry that significantly increased with age. It should also be noted that before FDR correction, as much as 15% of the bundles overall showed an increase in asymmetry with age (Figure 4E). Additional volume and streamline length results can be found in Supplementary figures 4 & 5.

#### Bundles showing accelerated decline with age in APOE4 carriers

We found no evidence of significant differences between APOE3 and APOE4 when examining the correlation between bundle metrics and age. Such differences could only be observed in 3 sub-bundles after splitting the subjects into three distinct groups of young, middle and old-aged individuals. These results can be seen in Figure 5 and Table 5 and correspond to a difference in genotype for average FA connecting insular sub-regions and the posterior parietal cortex for older individuals (lowered FA for APOE4, higher FA for APOE4), Figure 5.A, a difference in genotype for average FA near the posterior fusiform for young individuals (FA decreases for APOE3, increases for APOE4), Figure 5.B, and a difference in genotype for asymmetry of average FA of bundles between the superior frontal and the rostral middle frontal cortex, with the superior frontal region for older individuals (FA increases for APOE3, decreases for APOE4), Figure 5C.

**Figure 5.**
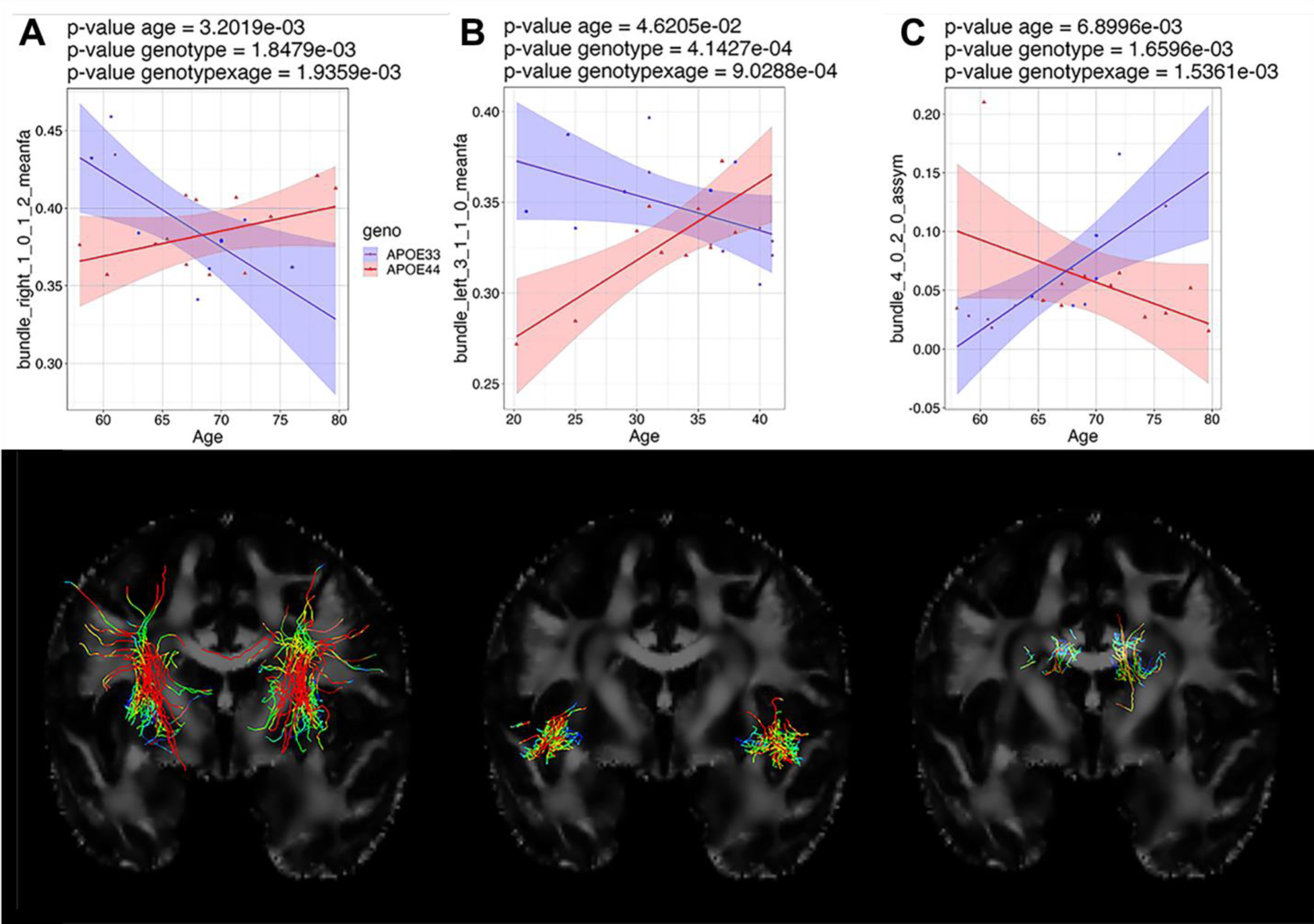
Genotype by age interactions deemed significant were found for both older and younger individuals.

**Table 5:**
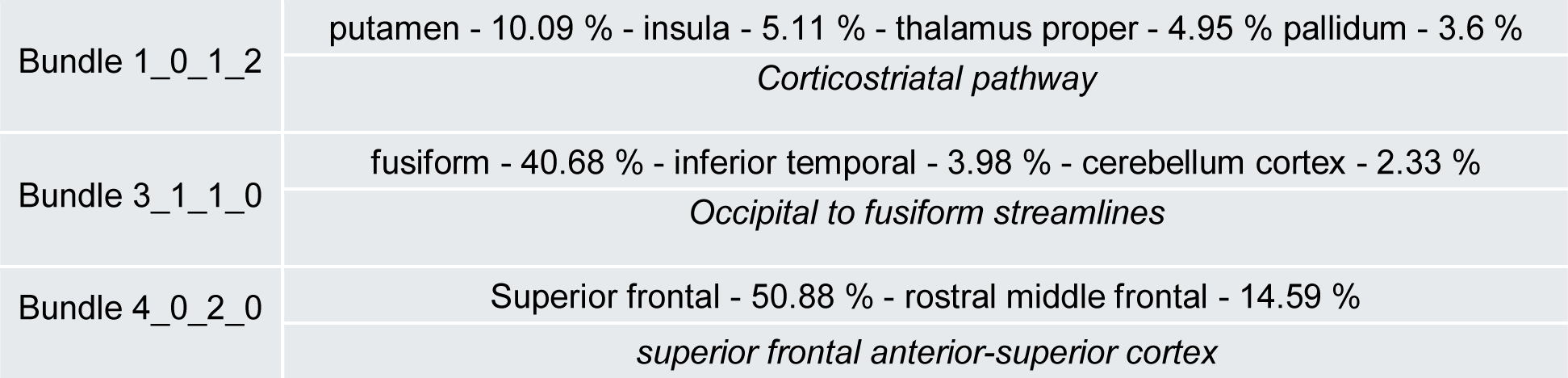
Correspondence between Bundle ID and top associated structures for bundles showing high correlation in differences of relationship between average FA or FA asymmetry and age.

### Cognitive traits association with physical and imaging metrics

#### Cognitive traits associated with aging

When focusing on the interaction between age and cognitive effects, we found that the greatest effect of age on cognition measures was connected to Visual attention. While it did not survive FDR correction, the correlation between Olfactory Memory and age behaved significantly differently between APOE3 individuals and APOE4 individuals, with APOE3 individuals having a stable or even increased olfactory memory over time, and the APOE4 individuals having an important decrease over time (Figure 6). Individual tests such as the useful field of view 2nd and 3rd, and the number of Digits for the Digit span, also showed a significant decrease even when considering all individual scores. Overall, the radar plots indicate worsening skills for most cognitive metrics when observing older individuals.

**Figure 6:**
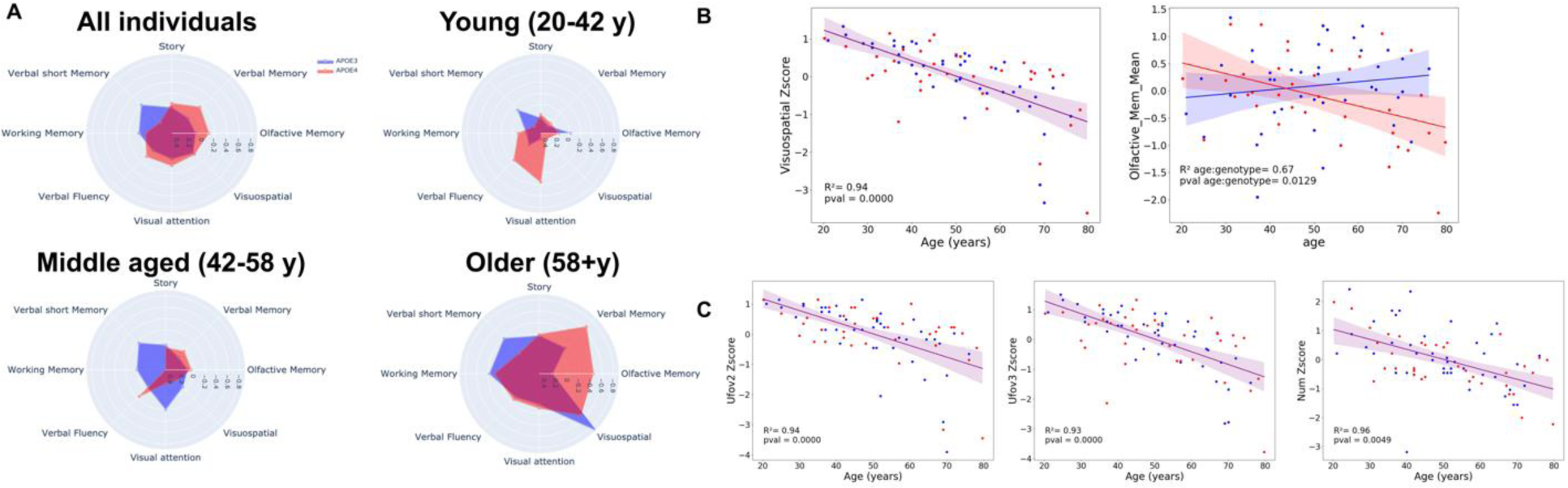
Cognitive changes indicated qualitative differences in visuospatial function, and olfactory memory between APOE genotypes (A). The quantitative analyses showed significance for the linear models for visuospatial function, olfactory memory, spatial attention, and digit symbol recognition, with a genotype specific pattern for olfactory memory (B).

**Table 6:** Correspondence between Bundle ID and top associated structures for bundles showing significant differences between genotypes for their correlation between the variance of FA with olfactory memory scores.

|  |  |
| --- | --- |
| Bundle 2 | Inferior parietal - 17.55 % lateral occipital - 10.74 % superior parietal - 9.56 % |
|  | *Parietal lobe* |
| Bundle 2_0 | Lateral occipital - 31.0 % pericalcarine - 12.75 % lingual - 11.02 % |
| Bundle 2_1 | Superior parietal - 25.81 % inferior parietal - 20.72 % precuneus - 12.64 % |
| Bundle 2_2 | Inferior parietal - 21.68 % lateral occipital - 5.15 % Bank of Superior Temporal Sulcus 4.83 % |
| Bundle 2_0_2 | Lingual - 28.76 % pericalcarine - 23.69 % fusiform - 1.28 % |
|  | *lingual to pericalcarine local connections* |
| Bundle 2_1_0 | Inferior parietal - 44.38 % superior parietal - 31.89 % precuneus - 4.16 % |
|  | *Lateral precuneus* |
| Bundle 2_1_1 | Superior parietal - 36.16 % cuneus - 16.0 % precuneus - 8.87 % |
|  | *Higher precuneus, lower cuneus* |
| Bundle 2_1_2 | Precuneus - 23.53 % cuneus - 5.42 % isthmus cingulate - 2.58 % |
|  | *Interhemispheric precuneus* |
| Bundle 2_2_0 | Inferior parietal - 32.2 % Bank of Superior Temporal Sulcus - 10.61 % |
|  | *Arcuate fasciculus superior* |
| Bundle 2_2_1 | Inferior parietal - 30.04 % lateral occipital - 17.09 % middle temporal - 5.23 % |
|  | *Arcuate fasciculus inferior* |
| Bundle 2_2_2 | Lingual - 7.94 % precuneus - 2.26 % pericalcarine - 1.16 % |
|  | *Interhemispheric cuneus* |
| Bundle 2_2_1_1 | Lateral occipital - 30.0 % middle temporal - 18.02 % inferior parietal - 10.54 % fusiform - 10.16 % |

#### Cognitive traits associated with APOE genotype and standard deviation of FA found in bundles

As previously seen in Figure 6, correlations between the Olfactory Memory and certain metrics are highly dependent on genotype and are even more so when considering specific bundles. A high standard deviation of FA across streamlines found in bundles associated with the posterior parietal and superior occipital lobe tends to lead to a higher olfactory memory score in APOE4 individuals, while it leads to a marginally lower score for APOE3 individuals. This is particularly true for those streamlines near the border of the parietal and occipital lobe, at the inferior parietal and lateral occipital cortices, as well as streamlines connecting lingual regions to the parietal and occipital lobe (Figure 7, Table 7)

**Figure 7:**
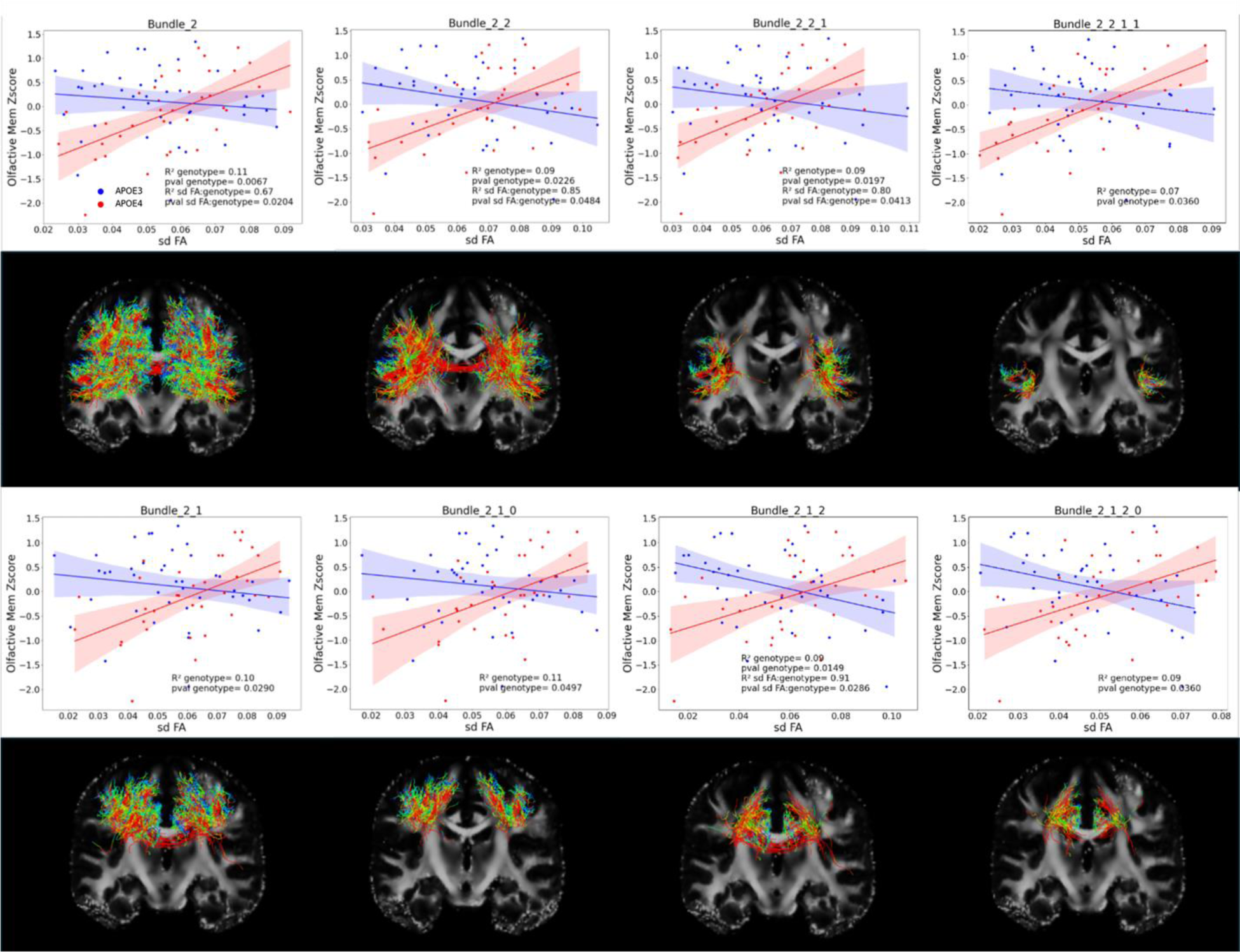
Cognitive scores genotype-specific significant interaction with variance of FA in parietal and lingual bundles

**Table 7:** Correspondence between Bundle ID and top associated structures, for bundles showing significant differences with genotype for their correlation between the volume of bundle streamlines or BUAN, and olfactory memory scores.

|  |  |
| --- | --- |
| Bundle 3_0_0 | Cerebellum cortex - 85.89 % |
| Bundle 3_0_1 | Cerebellum cortex - 44.81 % |
| Bundle 3_1_1 | Fusiform - 20.74 % lingual - 4.6 % hippocampus - 2.73 % |
| Bundle 4_2_0 | Rostral middle frontal - 25.77 % medial orbitofrontal - 6.29 % superior frontal - 3.33 % |
| Bundle 2 | Inferior parietal - 17.55 % lateral occipital - 10.74 % superior parietal - 9.56 % |
| Bundle 2_2 | Inferior parietal - 21.68 % lateral occipital - 5.15 % Bank of Superior Temporal Sulcus - 4.83 % |
| Bundle 2_2_1 | Inferior parietal - 30.04 % lateral occipital - 17.09 % middle temporal - 5.23 % |
| Bundle 2_2_2 | Lateral occipital - 2.72 %; inferior parietal - 2.68 %; fusiform - 2.12 % |
| Bundle 3 | Cerebellum cortex - 28.22 % fusiform - 3.88 % hippocampus - 3.85 % |
| Bundle 3_1 | fusiform - 9.4 % Bank Superior Temporal Sulcus - 6.97 % inferior temporal - 5.69 % |
| Bundle 3_1_2 | fusiform - 5.65 % hippocampus - 4.02 % superior temporal - 1.25 % |

#### Cognitive traits associated with APOE genotype and shape of bundles

The intersection between cognitive scoring and bundle shape metrics, specifically the volume of bundles and the BUAN between the left and right side of the bundles, is delineated in Figure 8.

**Figure 8:**
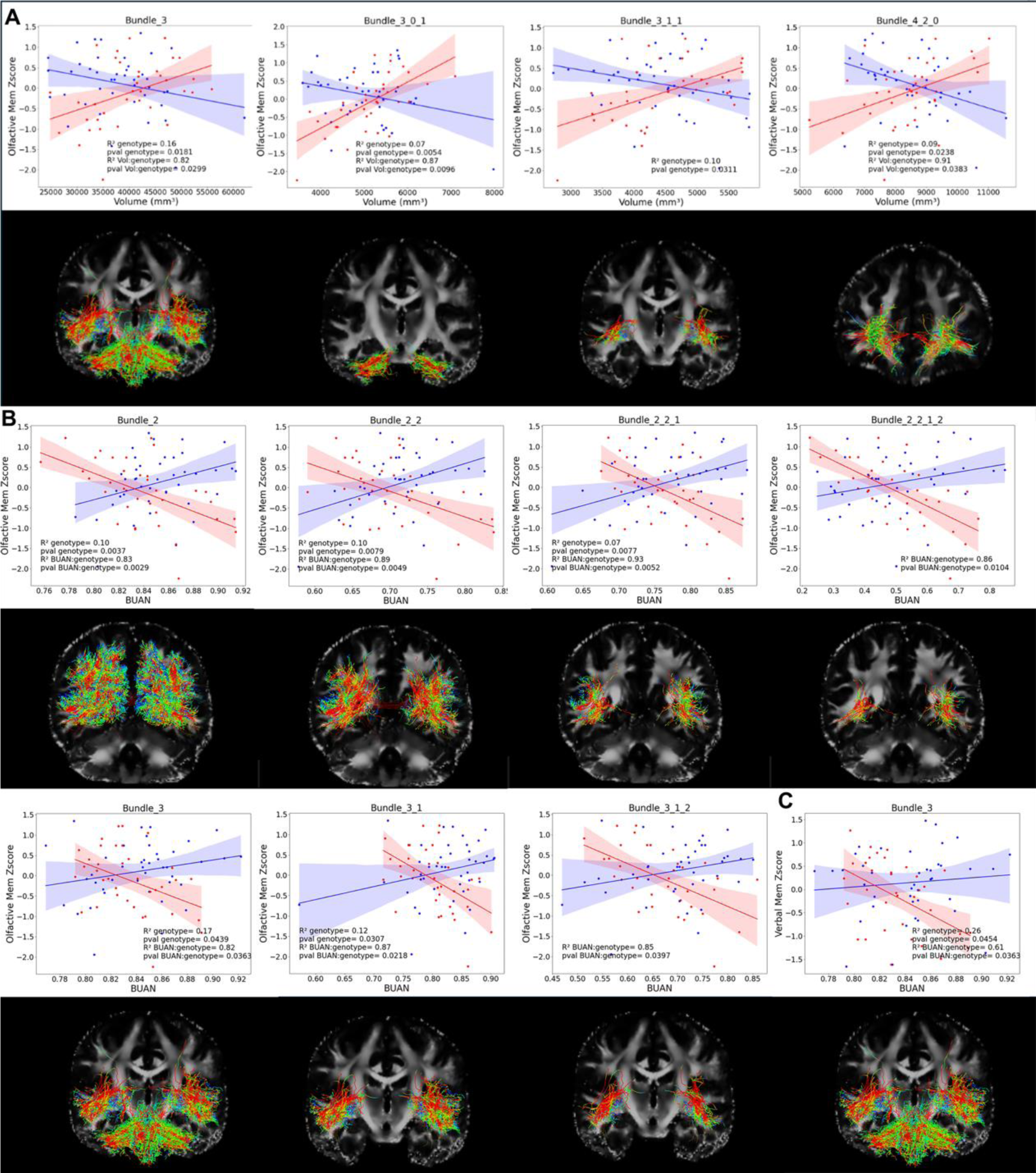
Genotype-specific significant interactions between cognitive scores and bundle metrics A) Olfactory score and bundle volume; B) Olfactory score and bundle similarity; C) Verbal Memory score and bundle similarity

Similarly to the results for the variance of FA, Olfactory Memory score was the only average cognitive score to display significant differences, with the behavior of bundle metrics versus Olfactory memory being highly dependent on the genotype of individuals. The volume of streamlines is one potential genotype differentiator, with a higher volume of certain bundles being associated with high olfactory score results in APOE4, while being associated with a lower cognitive score in APOE3 individuals. The bundles displaying this volume discrepancy are those found in the cerebellum, those connecting the occipital lobe with lingual regions, and streamlines found in the anterior frontal lobe (Figure 8A, Table 8).

**Table 8:** Correspondence between Bundle ID and top associated structures for bundles showing significant differences between physical traits of individuals and bundle statistics.

|  |  |
| --- | --- |
| Bundle 4_2_0 | Rostral middle frontal - 25.77 % medial orbitofrontal - 6.29 % superior frontal - 3.33 % |
| Bundle 4_2_0_1 | medial orbitofrontal - 23.06 % rostral middle frontal - 15.77 % frontal pole - 9.35 % |
| Bundle 5_2_0_0 | Rostral middle frontal - 21.74 % caudal middle frontal - 18.44 % superior frontal - 11.64 % |
| Bundle 4_1_1_1 | Pars triangularis - 44.98 % pars opercularis - 6.65 % rostral middle frontal - 3.79 % |
| Bundle 4_2_2 | Rostral anterior cingulate - 8.3 % superior frontal - 5.46 % medial orbitofrontal - 3.6 % |
| Bundle 5_2_1_0 | Superior frontal - 4.72 % rostral middle frontal - 3.37 % caudal middle frontal - 1.71 % |

BUAN also showed a similar genotype dependent differentiation in its correlation with Olfactory memory scores, and we observed that an increase of the BUAN metric would lead to an increase of the olfactory score in APOE3, while a highly symmetrical bundle in APOE4 is instead correlated with a lower olfactory score. Those bundles showing this genotype dependence on effect tended to be the same or neighboring the bundle streamlines previously identified by both FA standard deviation and the volume of streamline metrics, specifically those bundles associated with inferior parietal and lateral occipital regions as well as the cerebellum cortex (Figure 8B, Table 8). Additionally, we found that the verbal memory score and BUAN of cerebellum streamlines had a similar correlation as that found between the BUAN and the Olfactory memory score. (Figure 8C)

In summary, for the specific bundles found in the cerebellum, cuneus and inferior pre-cuneus, high left-right symmetry in APOE3 led to higher scores, high left-right symmetry in APOE4 led to lower olfactory score, high volume of streamlines led to slightly lower olfactory memory score for APOE3, high volume of streamlines led to higher scores for APOE4.

### Physiological traits associated with bundle statistics

Bundle metrics were compared with physiological traits of subjects and revealed that the average FA of the inferior frontal pole streamlines decreased with diastolic pressure (Figure 9.A, 9.B). Also, the FA of the streamlines of the anterior superior frontal cortex decreased with diastolic pressure (Figure 9.C). The average FA was also correlated with subject Height for the lateral frontal bundle streamlines of the brain. (Figure 9.D). The number of streamlines decreases with Systolic pressure for the bundle of streamlines responsible for interhemispheric connections of the left frontal lobe to the right frontal lobe (Figure 9.E).

**Figure 9:**
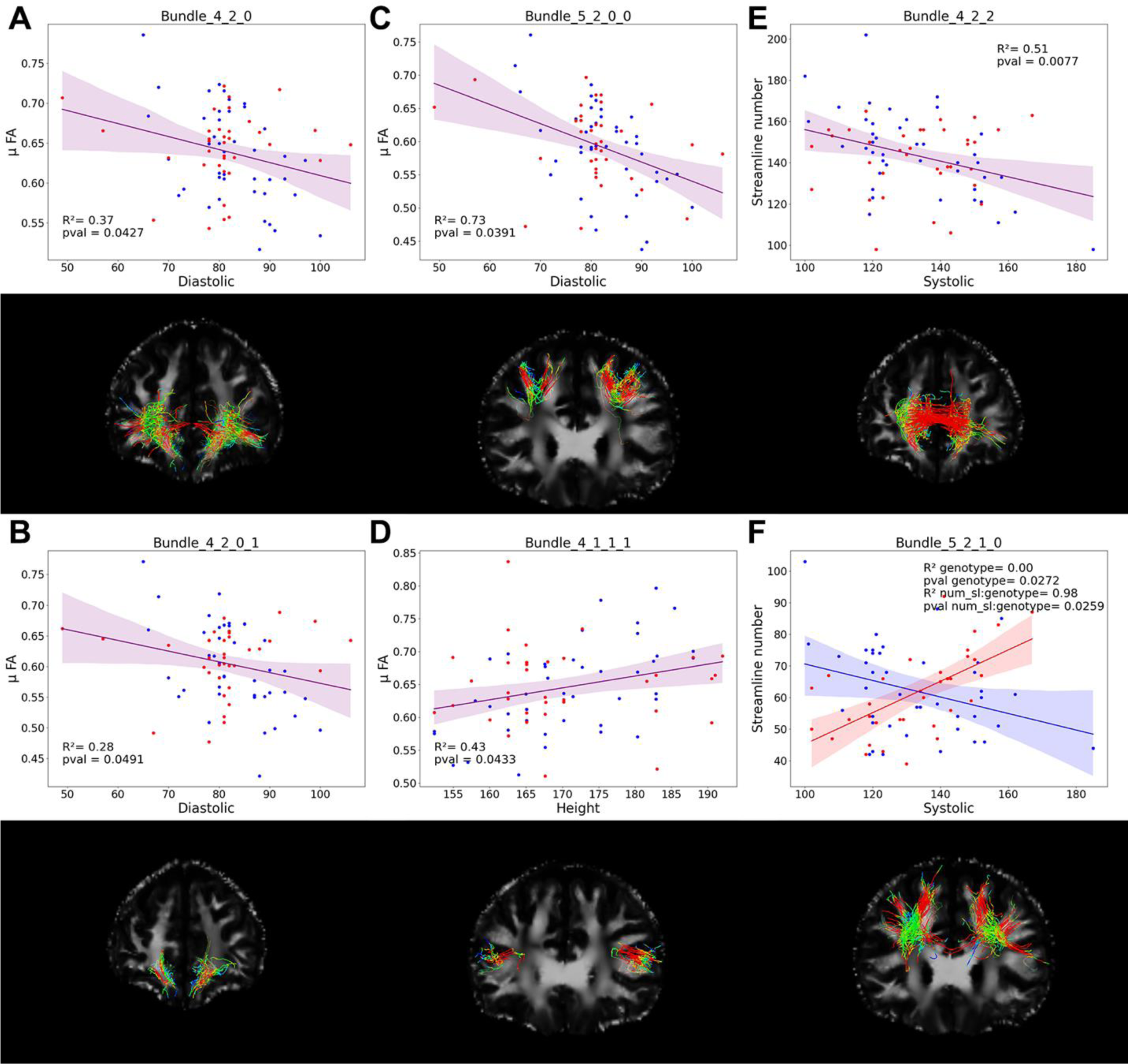
Significant bundle property changes due to physiological properties between: A, B, C) Diastolic pressure and average FA; D) Height and average FA; E) Systolic pressure and number of streamlines; F) Systolic pressure and genotype specific number of streamlines

Finally, there was one case of genotype differentiation, where the number of streamlines associated with the anterior superior frontal sub-cortical streamlines decreased with Systolic pressure in non-APOE4 carriers but increased in APOE4 individuals. (Figure 9.F)

## Discussion

Using a multi-scale analytical approach, we found that the FA of frontal lobe streamlines were generally the most affected by age, with important reductions in both average and standard deviation of FA found all across the frontal lobe streamlines, with this important FA decrease in the frontal lobe being corroborated by previous studies [51, 52].

However, this effect does not affect all frontal streamlines evenly, with streamlines associated with the rostral middle and superior frontal regions seeing greater age-related atrophy in average FA. While most reports concerning the rostral frontal gyrus are usually associated with schizophrenia [53, 54] or stress [55], changes in tracts local to that region as well as the superior frontal gyrus have also been associated with executive dysfunction in individuals with mild cognitive impairment (MCI) [56].

The superior frontal gyrus is known to be associated with both late life depression [57, 58] and most importantly, local microstructural degeneration is associated with early AD. [59]. The observed degeneration in test subjects is therefore a possible biomarker of developing AD, while our bundle splitting allows us to more precisely determine that this degeneration is particularly associated with a specific bundle within the frontal lobe, specifically those streamlines connecting the genu of the corpus callosum with the anterior frontal lobe and the corticostriatal tract (See Appendix Figure 1). A previous study of interest found that the FA of the corpus callosum genu was a biomarker for cognitive decay in older individuals [60].

We have reported that the inferior and middle longitudinal fasciculus saw a decrease in overall bundle volume. While there have been few studies concerning the evolution of the longitudinal fasciculus tracts with age, the reduction in volume of these tracts could potentially be related to the white matter cognitive impairment associated with these tract structures [61].

Our detected decrease in streamline length, volume and streamline number found in the superior parietal posterior cortex, meanwhile, is quite possibly linked to the observed cortical thinning of the parietal posterior cortex with age [62]. It should also be noted that this cortex also plays a compensatory role for atrophy in prefrontal areas [63].

When delving into the cognitive results of our study, we saw that age had a major effect on visuo-spatial ability. We also saw that the evolution of olfactory memory of individuals based on their age and phenotype was highly genotype dependent, with a major inflexion point in their early 40s. Further studies might elucidate that individuals with a faster degrading olfactory memory may develop AD, which has also been suspected by multiple recent studies [64–67]. Additionally, lack of APOE and the APOE4 allele itself has been linked to olfactory memory deficiencies, though the effects have almost only been observed in mice and olfactory neurites as of yet [67–71].

**In this** study, we compared these cognitive results with our bundle multi-level analysis and have found that the comparison of olfactory memory to specific metrics will show different behaviors depending on genotype in specific regions. Specifically, the two regions saw genotypical differentiation when compared with APOE.

First, APOE4 individuals that showed low FA variance and high shape symmetry in both the inferior precuneus and superior cuneus had low olfactory scores when compared to APOE3. Recent studies have both found that functional connectivity in the precuneus to the amygdala and hippocampus was linked to olfactory recognition [72] and that precuneus atrophy was linked to Alzheimer’s disease [73].

In the case of the cerebellum, we find that low tract volume and high shape symmetry lead to low olfactory memory scores in APOE4 compared to APOE3. While cerebellum is not traditionally linked to olfactory memory, some studies imply that it does play a role in overall sensory analysis along with other sensory regions [74–76]. Additionally, the cerebellum and olfactory ability are particularly linked when considering neurodegenerative diseases, such as Parkinson’s [77] and memory decline [78, 79].

In short, we have found the precuneus, cuneus, and the cerebellum symmetry to have genotype-specific interactions with olfactory ability in individuals, which could additionally be linked to functional study findings which show that the interaction of the cuneus and the cerebellum are related to olfactory ability [80]. These findings and our own results seem to indicate that the cerebellum and cuneus play a compensatory role in individual neurodegenerative risk, and that low tract spread and asymmetry in at risk individuals drastically affects olfactory ability and may foretell heightened risk of Alzheimer’s disease.

To summarize, in this study, we applied a multiscale approach to APOE and preclinical AD, which allowed us to find significant results which had previously eluded us, as observing tracts based on high impact connections using IIT-connectomes and TN-PCA or using Tract Seg did not lead to significant results. This may indicate that complex tract analysis could be superior to connectomic network organization where AD is concerned [81]. Our multiscale analysis allowed us not only to determine that the frontal lobe tract FA values were the most highly correlated with age, but also allow us to determine which specific subsets saw the greatest effect, while also finding genotype specific interactions between olfactory ability and the inferior precuneus, cerebellum and cerebellum tractography.

Limitations include that we only had one shell of b-values when acquiring the diffusion imaging data, and that our data set is still limited.

Further studies should more specifically address the streamlines connecting the genu of corpus callosum and the anterior frontal lobes, and their connections to the thalamus to determine relationships with genotype, and AD family history.

## Conclusion

While brain regional changes have taken center stage in examining the progression of AD, we propose a data driven approach to detect changes in populations at risk based on long range brain connections. Our hierarchical representation provides a sensitive approach to bundles that indicate through texture, variability, shape, and microstructural asymmetry, their specific vulnerability to risk factors for LOAD, e.g., age, APOE genotype. Further studies can investigate longitudinal aspects and the impact of modifiable risk factors.

## Acknowledgment

We are grateful for NIH support through RF1 AG057895, R01 AG066184, RF1 AG070149, and the Duke School of Medicine.

## Consent Statement

This study was approved by the Duke Institutional Review Board, and all subjects provided informed consent.

