## Supplementary material for "Mapping the impact of age and APOE risk factors for late onset Alzheimer’s disease on long range brain connections through multiscale bundle analysis": Supplental Figures

### Supplementary figures

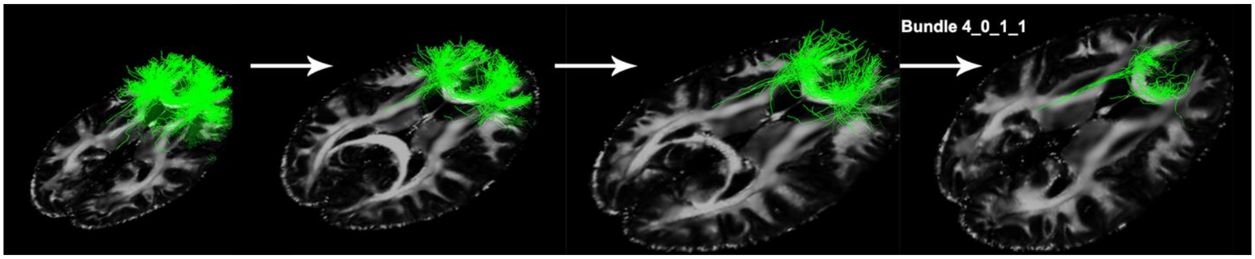


Supplementary Figure 1: Splitting of bundle 4, to bundle 4_0, 4_0_1, and ultimately 4_0_1_1

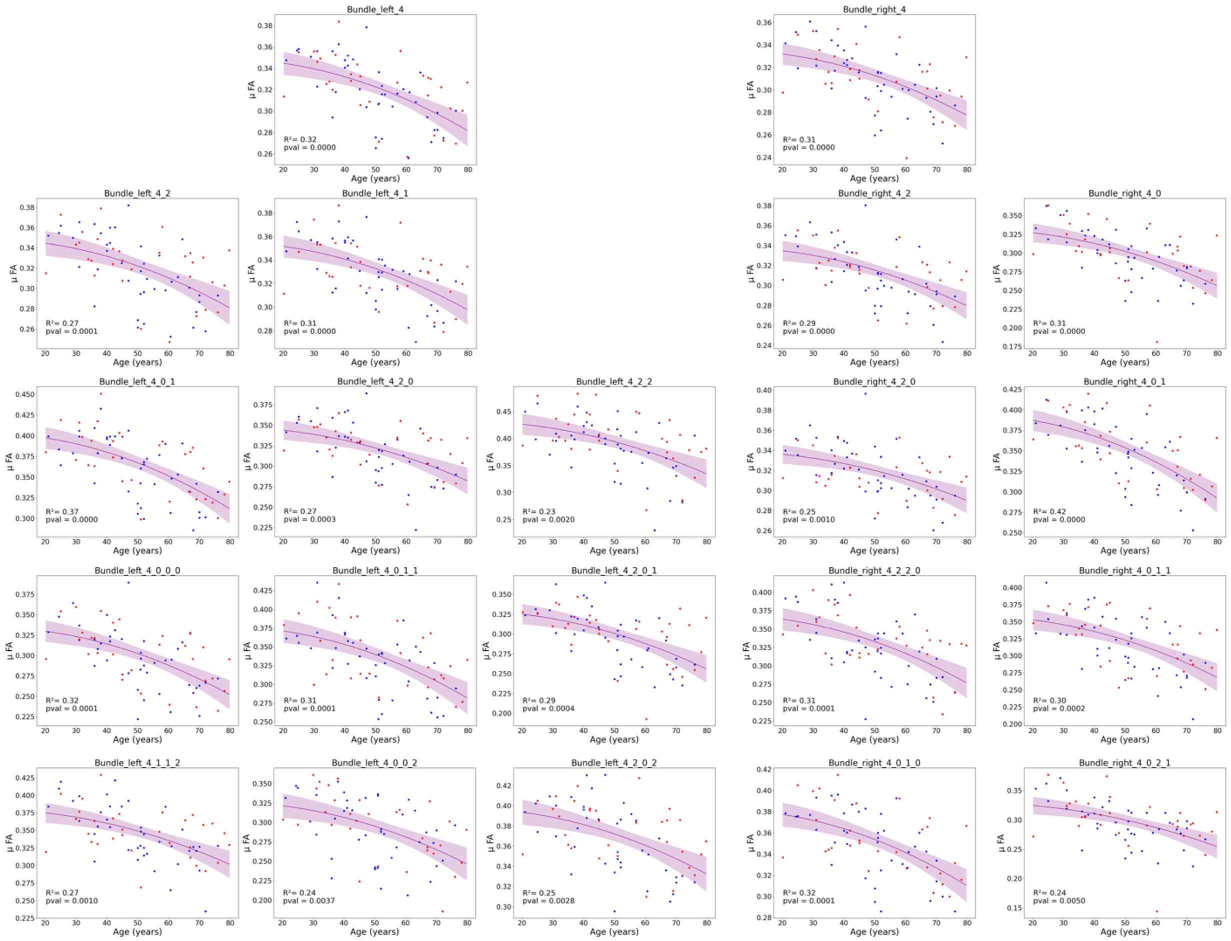


Supplementary Figure 2: Additional measurements for frontal lobe data of significant mean FA changes with age

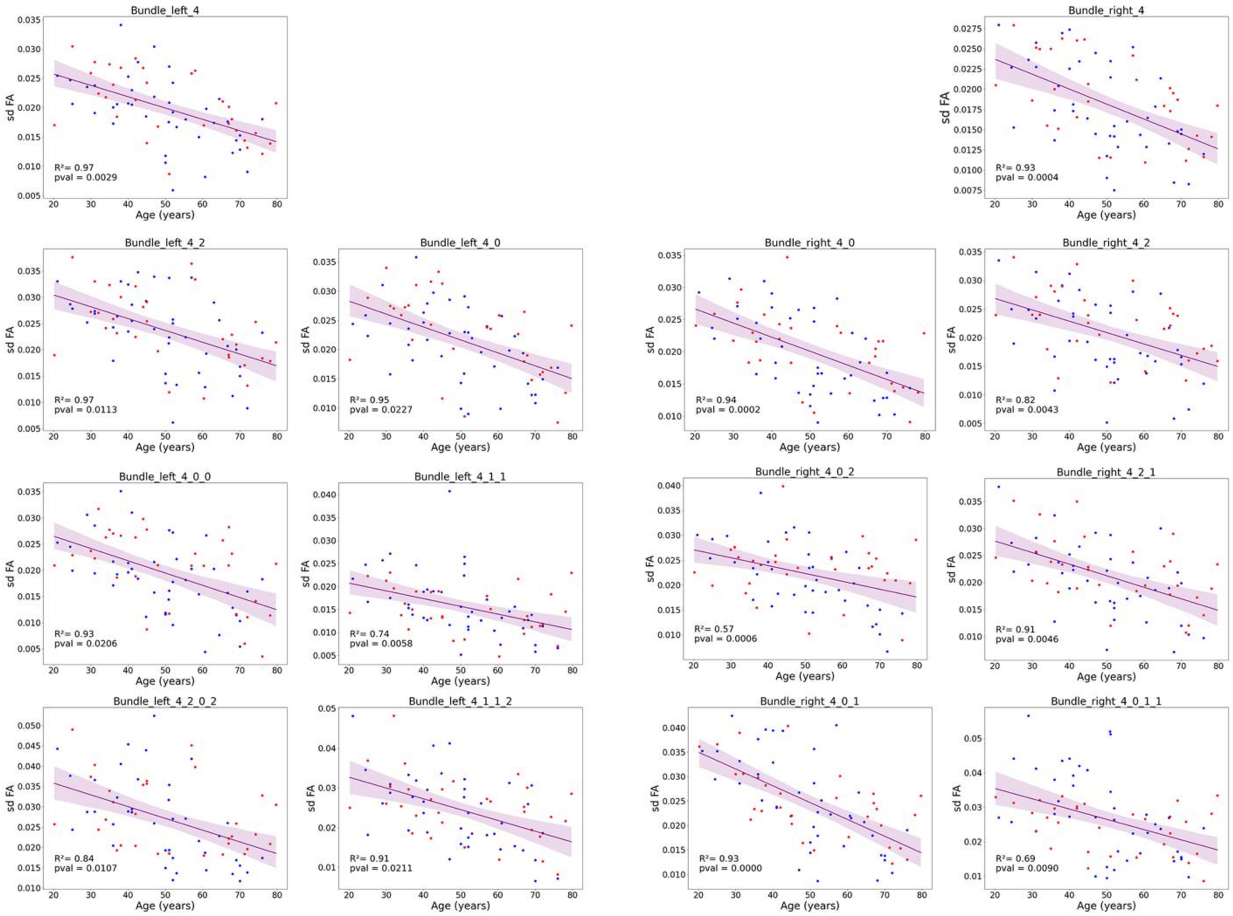


Supplementary Figure 3: Additional measurements for frontal lobe data of significant variance FA changes with age

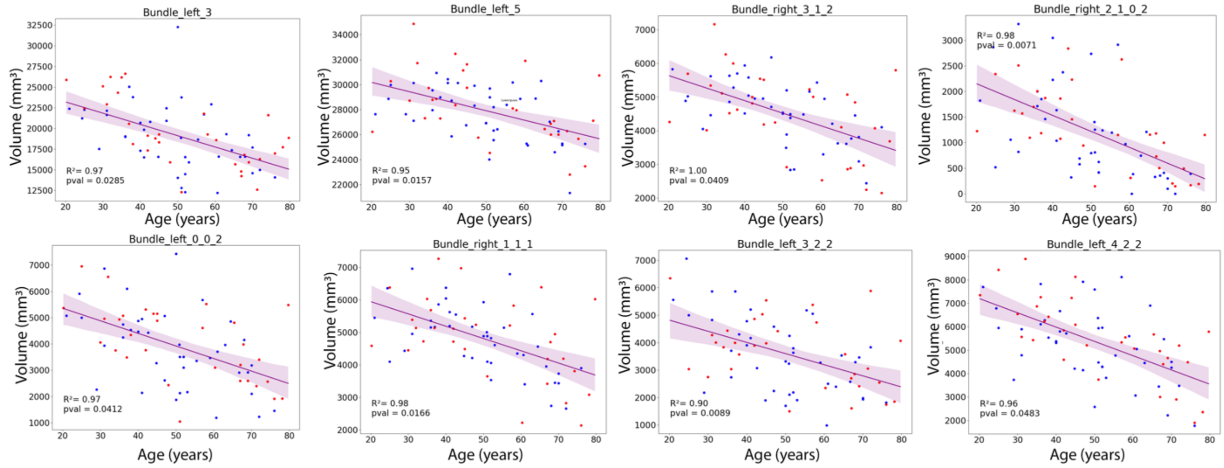


Supplementary Figure 4: Additional measurements for significant volume changes of bundles with age

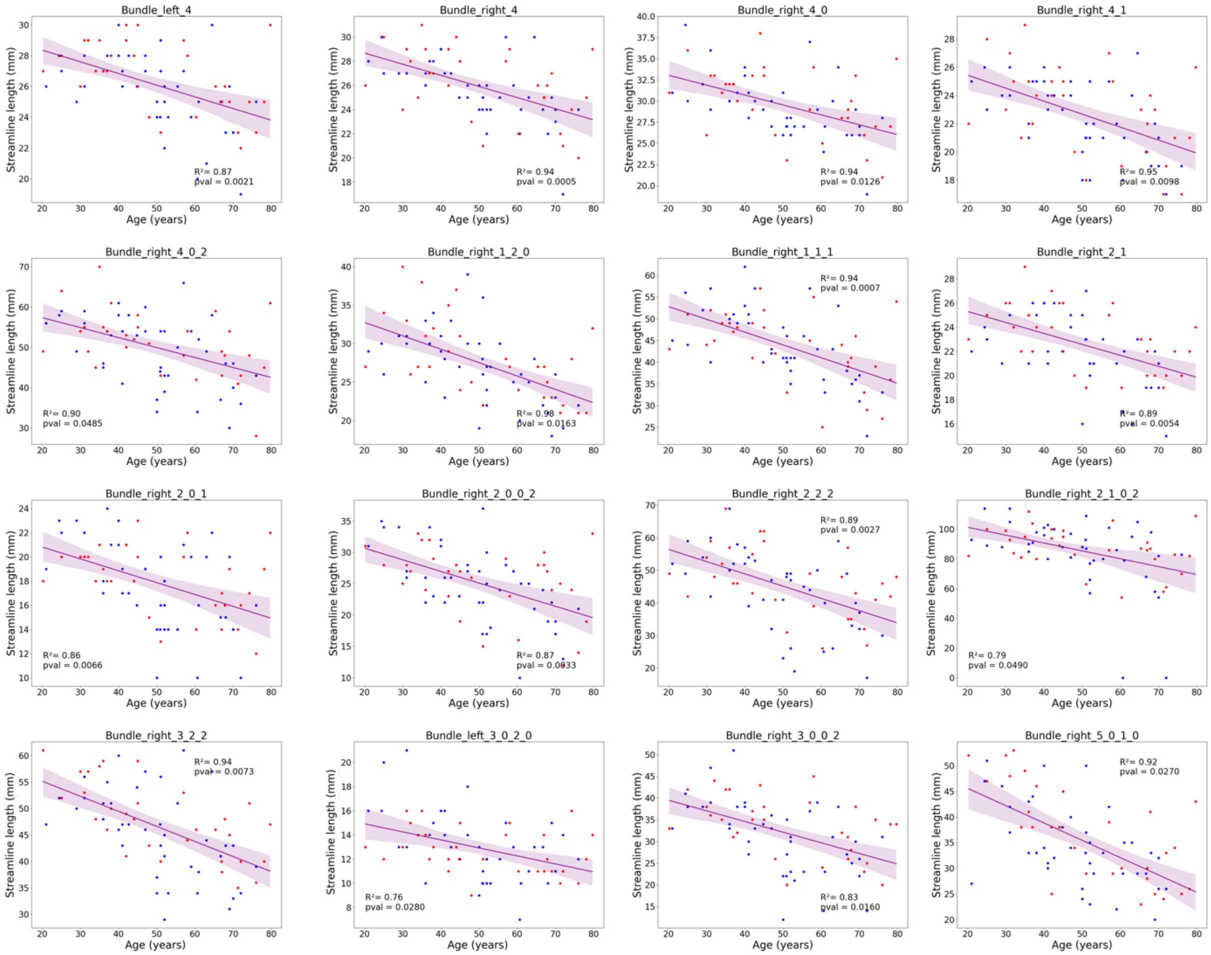


Supplementary Figure 5: Additional measurements for significant average streamline length changes with age
